# Antimicrobial peptides originating from expression libraries of *Aurelia aurita* and *Mnemiopsis leidyi* prevent biofilm formation of opportunistic pathogens

**DOI:** 10.1101/2023.03.02.530746

**Authors:** Lisa Ladewig, Leon Gloy, Daniela Langfeldt, Nicole Pinnow, Nancy Weiland-Bräuer, Ruth A. Schmitz

**Author notes:** Correspondence; (Germany). Joint first authorship (Lisa Ladewig, Leon Gloy).

## Abstract

The demand for novel antimicrobial compounds is rapidly growing due to the rising appearance of antibiotic resistance in bacteria; accordingly, alternative approaches are urgently needed. Antimicrobial peptides (AMPs) are promising since they are a naturally occurring part of the innate immune system and display remarkable broad-spectrum activity and high selectivity against various microbes. Marine invertebrates are a primary resource of natural AMPs. Consequently, cDNA expression (EST) libraries from the Cnidarian moon jellyfish *Aurelia aurita* and the Ctenophore comb jelly *Mnemiopsis leidyi* were constructed in *Escherichia coli*. Cell-free size-fractionated cell extracts (< 3 kDa) of the two libraries (each with 29,952 clones) were consecutively screened for peptides preventing the biofilm formation of opportunistic pathogens using the crystal violet assay. The 3 kDa fraction of ten individual clones demonstrated promising biofilm-preventing activities against *Klebsiella oxytoca* and *Staphylococcus epidermidis*. Sequencing the respective activity-conferring inserts allowed the identification of small ORFs encoding peptides (10 – 22 aa), which were subsequently chemically synthesized to validate their inhibitory potential. Biofilm-preventing effects against *K. oxytoca, Pseudomonas aeruginosa, S. epidermidis*, and *S. aureus* were verified for five synthetic peptides in a concentration-dependent manner, with peptide BiP_Aa_5 showing the strongest effects. The impact of BiP_Aa_2, BiP_Aa_5, and BiP_Aa_6 on dynamic biofilm formation of *K. oxytoca* was further validated in microfluidic flow cells, demonstrating a significant reduction in biofilm thickness and volume by BiP_Aa_2 and BiP_Aa_5. Overall, the structural characteristics of the marine invertebrate-derived AMPs, their physicochemical properties, and promising anti-biofilm effects highlight them as attractive candidates for discovering new antimicrobials.

## 1. Introduction

Multicellular organisms evolved in the presence of microbes, which play fundamental roles in their health, development, and evolution [1]. Early-branching metazoans, i.a., invertebrates, have required complex systems for recognizing and discriminating beneficial and pathogenic microorganisms [2-5]. However, unlike vertebrates, invertebrates lack classical antibody-based adaptive immunity [2]. Cellular immunity in invertebrates originates from defense reactions (innate immune system), including hemocyte-mediated modules, encapsulation, phagocytosis, and generation of antimicrobial peptides (AMPs) [6]. AMPs are small, mainly positively charged peptides that promote the innate defense mechanism by targeting the negatively charged membranes of microorganisms [7]. AMPs get embedded in the hydrophobic regions of lipid membranes, often forming pores, thus destabilizing biological membranes and causing cell lysis [8,9]. They are rapidly induced in response to microbes to modulate the immunoreactions. AMPs have a wide range of inhibitory effects against bacteria, fungi, parasites, and viruses [10]. Consequently, AMPs have awakened interest as potential next-generation antibiotics since emerging antibiotic resistance in pathogenic bacteria is a serious challenge and has led to the need for new alternative bioactive molecules less prone to bacterial resistance [11,12]. A particular interest in using AMPs is to combat biofilms of pathogens that have been demonstrated to be significant contributors to diseases [13,14]. Multiple factors contribute to the overall resistance of biofilms against antibiotics, including reduced metabolic- and growth rates, protection by extracellular polymeric substances, and specific resistance mechanisms conferred by the altered physiology of biofilm bacteria [15,16]. Focusing on treating the highly challenging, adverse effects of harmful biofilm-associated infections, AMPs are indicated to have strong potential as antimicrobials and antibiofilm agents [13,17-19]. Marine invertebrates are the primary source of natural AMPs [6]. Over the past decades, several AMPs have been identified and isolated from marine invertebrates, including cnidarians, mollusks, annelids, arthropods, and tunicata [20]. Those peptides showed a broad spectrum of antimicrobial properties against Gram-negative and Gram-positive bacteria. They possess novel and unique structures by only exhibiting a few side effects [6].

The present study aimed to identify novel antimicrobial peptides from lower metazoans to prevent biofilms. Consequently, cDNA expression (EST) libraries were constructed from the Cnidarian moon jellyfish *Aurelia aurita* and the Ctenophore comb jelly *Mnemiopsis leidyi*. Low-molecular-weight fractions (< 3kDa) derived from both EST libraries were functionally screened to identify activities preventing the biofilm formation of opportunistic pathogens. After identifying the sORFs in the respective inserts, the corresponding peptides were chemically synthesized and tested for their potential to prevent static and dynamic pathogenic biofilms.

## 2. Materials and Methods

### 2.1. *Aurelia aurita* polyp husbandry

Husbandry of polyps is described in detail in previous studies by Weiland-Bräuer *et al*. [21,22]. Polyps of sub-population North Atlantic (Roscoff, France) were kept in the laboratory at 20 °C in 30 PSU artificial seawater (Tropical Sea Salts, Tropic Marin) and fed twice a week with freshly hatched *Artemia salina* (HOBBY, Grafschaft-Gelsdorf, Germany).

### 2.2. Sampling of *Aurelia aurita* polyps and *Mnemiopsis leidyi* medusae

*A. aurita* polyps were used from husbandry by removing individual polyps with a disposable pipette. Individual *M. leidyi* medusae (with a mean umbrella diameter of 4 cm) were sampled from the Kiel Bight, Baltic Sea (54°32.8’N, 10°14.7’E) in May 2017 using a dip net. The animals were immediately transported to the laboratory and washed thoroughly with sterile artificial seawater.

### 2.3. Direct mRNA isolation and construction of the cDNA expression libraries

The mRNA of *A. aurita* polyps and *M. leidyi* medusae were isolated with the DynaBeads® mRNA DIRECT Micro Kit (Ambion, Austin, USA) according to the manufacturer’s protocol “mRNA isolation from tissues”. In total, 28 parallel preparations were conducted using pools of 10 *A. aurita* polyps and 1×1 cm parts of *M. leidyi* medusae (in total 10 medusae) per isolation. Animal tissues were frozen in liquid nitrogen and homogenized with a motorized pestle (polyps) or a blender (medusae). For *A. aurita*, 3 μg and for *M. leidyi* 1.2 μg mRNA were used to construct cDNA expression libraries with the Clone MinerII cDNA Library Construction Kit (Invitrogen, Waltham, USA) according to the manufacturer protocol. First, mRNA was transcribed into cDNA and subsequently cloned into the pDONR222 entry vector. Second, the entry clones were recombined with Gateway expression vector pET300/NT-DEST using the ChampionTM pET300/NT-DEST GatewayTM Vector Kit (Invitrogen, Waltham, USA) to create expression clones. The resulting plasmids retained the original alignment and reading frame, allowing for functional analysis of full-length genes and entire libraries. SoluBL21 electrocompetent *Escherichia coli* cells were used for electroporation and as background strain for expression (Genlatis, San Diego, USA). Each library consisted of 29,952 single clones, stored in 96-well microtiter plates at -80 °C in the presence of 8 % DMSO as cryoprotectant.

### 2.4. Preparation of cell-free size-fractionated cell extracts

Single cDNA clones were grown in 200 μL Luria Bertani medium (LB, Carl Roth, Germany) supplemented with 100 μg/mL ampicillin overnight at 37 °C in 96-well microtiter plates. Initially, 96 clones were pooled. The mixture of bacterial cultures was centrifuged at 9,000 x g for 5 min, and the bacterial pellet was resuspended in 250 μL 50 mM Tris/HCl buffer (50 mM NaCl, pH 7.8). The suspension was transferred to screw-cap tubes (Sarstedt, Nümbrecht, Germany) containing one glass bead of the diameter 2.7 mm and approximately 50 mg of glass beads of the diameter 0.1 mm (Carl Roth, Karlsruhe, Germany). Snap-frozen (liquid N_2_) bacterial cells were mechanically disrupted with Precellys® (Bertin instruments, Montignyle-Bretonneux, France). The cell extract was sterile-filtered through a 0.2 μm centrifugal filter (Amchro GmbH, Hattersheim am Main, Germany) followed by size-fractionation using a 3 kDa centrifugal filter according to the manufacturer’s instructions (Merck KGaA, Darmstadt, Germany). The samples were kept at 4 °C or on ice throughout the process. The fractions < 3 kDa were collected, and the approximate protein concentrations were measured using a NanoDrop1000 (Thermo Fisher Scientific, Waltham, USA) and adjusted to the same concentration. The empty pET300/NT-DEST/*E. coli* SoluBL21 was used as a control. Biofilm-preventing 96-pools were further tested in pools of 48 and 24 clones to unravel single clone(s) responsible for the biofilm inhibition.

### 2.5. Bacterial biofilm prevention *in vitro* assay

Opportunistic pathogenic bacteria *K. oxytoca, P. aeruginosa* PAO1, *S. epidermidis* RP62A, and *S. aureus* were grown in 5 mL LB medium overnight at indicated temperatures (**Tab. 1**). Cell concentrations of overnight cultures were analyzed by Neubauer cell counting and set to 3 × 10^8^ cells/mL using GC minimal medium (with 1% (v/v) glycerol, 0.3% (w/v) casamino acids) [23] for *K. oxytoca* and Caso bouillon (17 g/L casein peptone, 3 g/L soybean peptone, 5 g/L NaCl, 2.5 g/L K2HPO4, 2.5 g/L glucose) for the remaining strains in the crystal violet assay. Cultures were aliquoted in 96-well plates (180/195 μL for each cavity). Cell-free size-fractionated cell extracts (20 μL) and synthesized peptides (various concentrations, 5 μL) were added to the cultures. MTPs were closed with gas-permeable membranes (Breathe-Easy®, Diversified Biotech, Dedham, USA) and incubated at 37 °C (*P. aeruginosa, S. aureus, S. epidermidis*) or 30 °C (*K. oxytoca*) for 18 h. Three biological were performed, each with eight technical replicates. Medium controls were performed for normalization. Biofilm formation was monitored and quantified using the crystal violet assay by measuring the absorbance of resolved crystal violet (70 % ethanol) at 590 nm with plate reader Spectra max Plus 384 (Molecular Devices, Ismaning; DE) [24,25]. The screening procedure is depicted in **Figure S1**. In addition, the potentially growth-inhibiting effects of synthetic peptides were tested on planktonic growing pathogens. Again, 3 × 10^8^ cells/mL diluted in LB medium were aliquoted in 96-well plates (200 μL for each cavity). Synthetic peptides were added to the final concentrations of 3.5 μg/mL and 112.5 μg/mL at the beginning of the experiment. MTPs were closed with gas-permeable membranes and gently shaken at 80 rpm and 37 °C (*P. aeruginosa, S. aureus, S. epidermidis*) or 30 °C (*K. oxytoca*) for 18 h. Turbidity at 600 nm was measured with plate reader Spectra max Plus 384.

**Table 1:** Bacterial strains used in this study.

| Bacterial strain | Description | Growth medium | Biofilm formation medium | Growth temperature [°C] |
| --- | --- | --- | --- | --- |
| *Pseudomonas aeruginosa* PAO 1 | DSM 1707 | LB | GC medium | 37 |
| *Staphylococcus aureus* | DSM 11823 | LB | Caso bouillon | 37 |
| *Staphylococcus epidermidis* RP62A | DSM 28319 | LB | Caso bouillon | 37 |
| *Klebsiella oxytoca* M5aI | DSM 7342 | LB | Caso bouillon | 30 |

### 2.6. Plasmid preparation, insert size determination, and sequencing

Plasmid DNA of identified biofilm-preventing single clones was isolated from 5 mL overnight cultures using the Presto™ Mini Plasmid Kit, according to the manufacturer (Geneaid, Taiwan). A restriction digest with the *Bsr*GI enzyme was performed for insert size determination, followed by subsequent gel analysis. The plasmids were Sanger-sequenced by the Institute of Clinical Molecular Biology, CAU Kiel, Germany. Sequences were analyzed with Geneious Prime software (Biomatters, Auckland, New Zealand). Vector sequences and PolyA tails were removed from the raw sequences. Cleaned insert sequences were used for open reading frame identification. The peptide sequences within the frame of the vector coding Histidine-tag were selected for peptide synthesis.

### 2.7. Synthesis of peptides

Peptides with sufficient biofilm-preventing activity and a length of > 5 amino acids were synthesized by conventional solid-phase peptide synthesis with > 94 % purity at Genscript (Leiden, Netherlands) (**Tab. 2**). The synthetic peptides were dissolved in sterile distilled water (Carl Roth, Karlsruhe, Germany) and stored at -80 °C as 100 μL aliquots of 10 mg/mL.

**Table 2:**
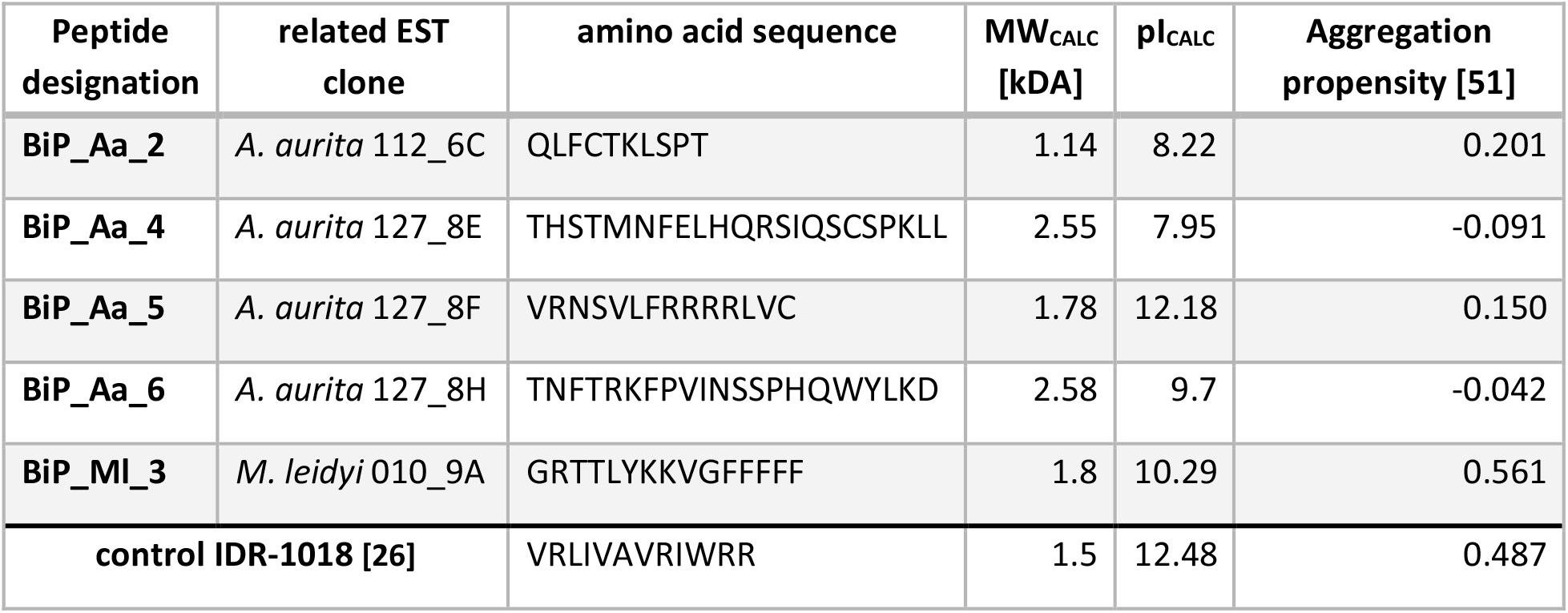
Synthetized peptides. Peptides were synthesized in 94 % purity at Genscript (Leiden, Netherlands). The characteristics of the peptides are summerized. MW_CALC_, calculated molecular weight; pI_CALC_, isoelectric point

### 2.8. Effect of biofilm-preventing peptides on biofilm formation of *K. oxytoca* M5aI in a microfluidic flow cell

The microfluidic flow cell system comprises a polymethyl methacrylate corpus with a single channel (1 × 8 × 0.1 mm). Two inlets and one outlet exist (**see Fig. 3A**). A borosilicate glass (24 × 60 × 0.17 mm; Carl Roth, Karlsruhe, Germany) was fixed with adhesive Black-Seal silicone (Weicon, Münster, Germany). Microliter syringes (Innovative Labor System GmbH, Stützenbach, Germany) were connected via Polytetrafluoroethylene (PTFE) tubes (0.3 mm x 0.6 mm) (Bola, Grünsfeld, Germany) with the in- and outlets and clamped into a syringe pump (Model: 220) (KD Scientific, Holliston, USA). The channel was sterilized by rinsing 70% EtOH for 24 h at a flow rate of 20 μL/h. Afterwards, the remaining ethanol was washed out with sterile water at 20 μL/h for 1 h. Subsequently, the channel was equilibrated with GC medium at 20 μL/h for 2 h. Inoculation was conducted with a cell suspension of 1 × 10^9^ cells/mL, generated from an overnight culture of *K. oxytoca* M5aI. Initial cell adhesion was ensured by incubation at 30 °C for 1 h without any flow. Next, the channel was continuously rinsed at 15 μL/h with GC medium injected within the first inlet. The synthetic peptides IDR-1018 (biofilm-preventing control peptide, [26,27], BiP_Aa_2, BiP_Aa_5, and BiP_Aa_6 were diluted in GC medium to a concentration of 25 μg/mL and injected within the second inlet. Thus, the final peptide concentrations were 12.5 μg/mL, corresponding to 10 ng peptide within the 0.8 μL channel. Biofilm formation was conducted at a flow rate of 15 μL/h, for 24 h and at 30 °C. Biofilm formation of *K. oxytoca* M5aI was further studied without adding peptides. Four biological replicates were conducted for each treatment.

Biofilms were stained with the fluorescent dye Syto9 (488 nm) (Life Technologie, Carlsbad, USA) and analyzed using the LSM 700 confocal laser scanning microscope (Zeiss, Oberkochen, Germany). Therefore, the microfluidic flow cells were washed with GC medium at 3 μL/min for 15 min. The channel was then filled with a 1:1000 dilution of the fluorescent dye Syto9 and incubated in the dark at room temperature for 30 min. Dye residues were removed by rinsing GC medium at 3 μL/min for 15 min. Four image stacks were recorded per flow cell with optical sections of 0.9 μm per z-step. The digital image acquisition, three-dimensional reconstruction, and calculation of the biofilm parameters were performed with the software “Zen Black” (Zeiss, Oberkochen, Germany) and the microscopy image analysis software “Imaris” (Oxford Instruments, Abingdon, UK).

## 3. Results

Expression libraries of two basal metazoans, *A. aurita* and *M. leidyi* were constructed and screened for biofilm-preventing peptides against opportunistic pathogenic bacteria (*K. oxytoca, P. aeruginosa, S. epidermidis, S. aureus)* using the crystal violet assay. Promising peptide candidates were characterized for their potential to prevent dynamic biofilm formation of *K. oxytoca* in microfluidic cells.

### 3.1 Construction of cDNA expression libraries of two basal metazoans

Expression libraries were constructed from mRNA derived from the Cnidarian moon jellyfish *A. aurita* and the Ctenophore comb jelly *M. leidyi* into the Gateway expression vector pET300/NT-DEST (Invitrogen). Each library consisted of 29,952 single clones (pET300/NT-DEST in *E. coli* Solu BL21). According to the manufacturer, the generated plasmids of the expression clones retained the original alignment and reading frame of the insert, allowing for functional analysis of full-length genes. Plasmids of randomly selected clones (96 clones per library) were characterized by restriction analysis, demonstrating an average insert size of approximately 1.4 kbps and 95 – 98 % insertion efficiencies for both libraries. Analyzing the insert sequences of those randomly selected clones revealed that 92 % of the reads aligned to the respective genomes [28–31].

### 3.2 Identification of biofilm-preventing clones from the cDNA expression libraries

Both constructed cDNA libraries were used to identify biofilm-preventing peptides derived from the basal metazoan hosts. Cell-free cell extracts were successively prepared from pools of 96, 48, 24, and single clones of the cDNA expression library (**Fig. S1**). Notably, cell extracts were size-fractionated using 3 kDa cutoff columns, and only the fractions < 3 kDa were analyzed using the crystal violet assay. This assay was initially performed with the Gram-negative opportunistic pathogen *K. oxytoca* and the Gram-positive pathogen *S. epidermidis*. In total, 36 % of the 96 clone pools showed biofilm-preventing activity for at least one of the two tested pathogens. Successively, pools of decreasing clone numbers were screened until single clones were identified (**Fig. S1**). Cell-free size-fractionated cell extracts of single clones were evaluated for their biofilm-preventing potential, including two additional opportunistic pathogens, *P. aeruginosa* and *S. aureus*. Overall, ten biofilm-preventing single clones (6 derived from *A. aurita*, and 4 from *M. leidyi*) were identified with varying impacts on biofilm formation of the different pathogens (**Tab. 3**). Sequence analysis of the biofilm inhibition-conferring inserts identified the corresponding small open reading frames (sORFs) (**Tab. S1**). Here, exclusively those sORFs N-terminally fused to the vector-derived Histidine-tag and longer than five amino acids were used for further analysis. Structural models of the five respective peptides were predicted with PEP-FOLD 3 (**Fig. 1**). NCBI BLASTp identified those peptide sequences in *A. aurita* and *M. leidyi* genomes. However, non-significant E-values were reached due to the short input sequences.

**Figure 1:**
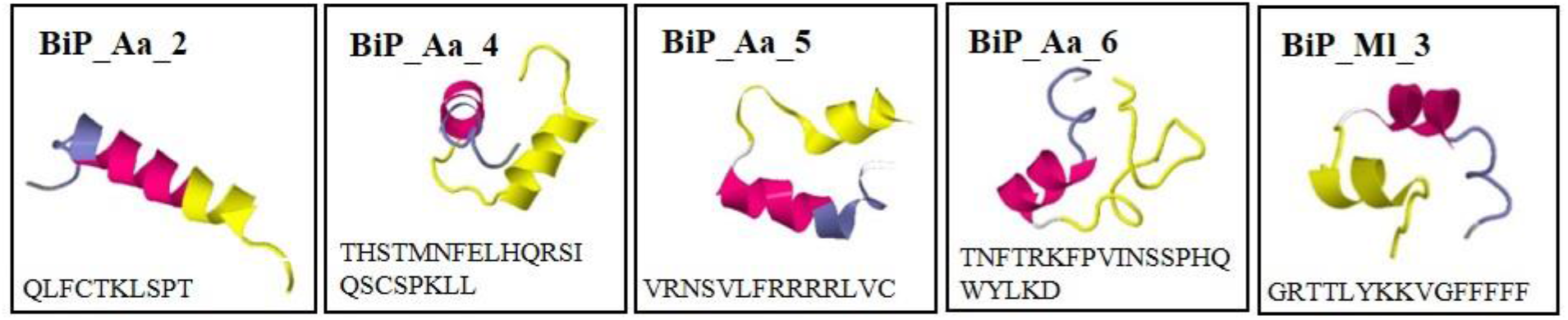
Predicted structural models of biofilm-preventing peptides. Sequences were analyzed using the Geneious Prime software (Biomatters, Auckland, New Zealand). Models were created using PEP-FOLD 3.

**Table 3:** Identified single clones derived from the cDNA expression libraries of *A. aurita* and *M. leidyi* and their impact on the biofilm formation of pathogens. Biofilm biomasses were calculated based on the crystal violet assay. Biofilm formation of pathogens without adding a test substance was set as 100 %.

| Clone designation | Biofilm formation [%] |  |  |  |
| --- | --- | --- | --- | --- |
|  | *K. oxytoca* | *P. aeruginosa* | *S. aureus* | *S. epidermidis* |
| Aa_112_4H | 71 ± 12 | 85 ± 19 | 77 ± 9 | 55 ± 14 |
| Aa_112_6C | 92 ± 14 | 79 ± 24 | 84 ± 13 | 49 ± 9 |
| Aa_127_8A | 22 ± 2 | 100 ± 18 | 100 ± 13 | 77 ± 13 |
| Aa_127_8E | 80 ± 3 | 69 ± 17 | 78 ± 13 | 6 ± 1 |
| Aa_127_8F | 28 ± 3 | 75 ± 18 | 84 ± 23 | 95 ± 10 |
| Aa_127_8H | 104 ± 5 | 77 ± 18 | 61 ± 10 | 40 ± 12 |
| MI_068_11H | 107 ± 13 | 81 ± 13 | 69 ± 7 | 29 ± 3 |
| MI_011_11H | 27 ± 2 | 100 ± 22 | 67 ± 3 | 77 ± 10 |
| MI_010_9A | 71 ± 8 | 78 ± 14 | 76 ± 5 | 94 ± 9 |
| MI_010_9G | 68 ± 14 | 75 ± 18 | 89 ± 17 | 91 ± 7 |

### 3.3 Synthetic peptides - verification of the biofilm-preventing effects on static biofilms

The five identified peptides were chemically synthesized and designated as BiP_Aa_2, BiP_Aa_4, BiP_Aa_5, BiP_Aa_6, and BiP_Ml_3 (**Tab. 2**). As a control, peptide IDR-1018 was additionally synthesized, which has been reported to inhibit planktonic bacteria [26,27]. Initially, potential growth-inhibiting effects of the synthetic peptides on planktonic growing pathogens were evaluated with two contrasting concentrations of 3.5 and 112.5 μg/mL. None of the peptides affected growth behaviour of any of the pathogens at the low concentration. However, in accordance with the original report [26,27] control peptide IDR-1018 administered in high concentrations significantly reduced planktonic growth of all strains (**Fig. 2**).

**Figure 2:**
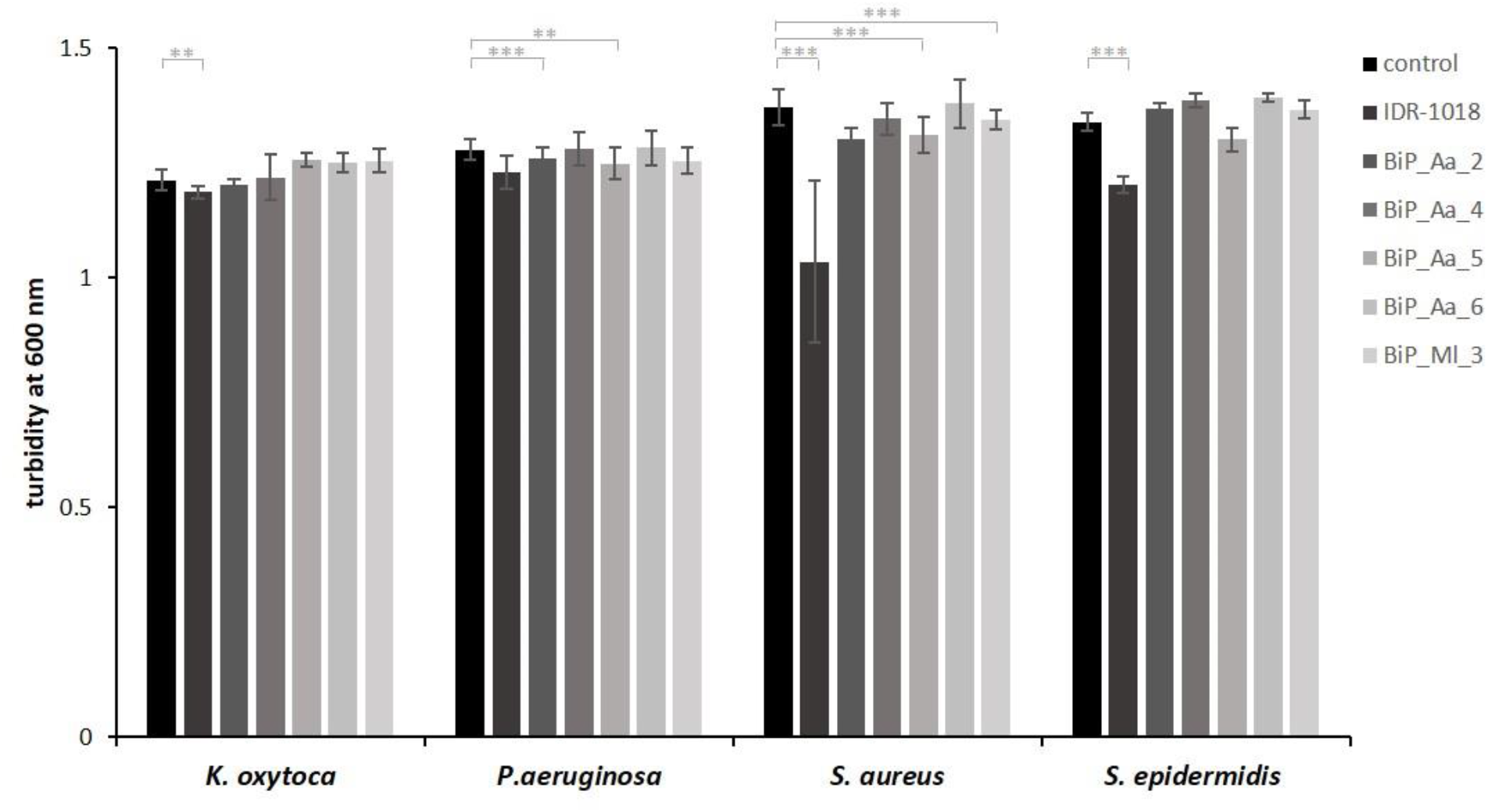
Exclusion of potential growth-inhibiting effects of the synthetic peptides on planktonic cells. *K. oxytoca, P. aeruginosa, S. epidermidis, and S. aureus* (3 × 10^8^ cells/mL) were grown for 18 h in 200 μL LB medium at 80 rpm after adding 112.5 μg/mL of the synthetic peptides. Turbidity was monitored at 600 nm of two biological, each with eight technical replicates, represented as means. The growth-inhibiting effects were compared to the control without addition of peptide. Significant p-values are indicated as P < 0.01,**, and P < 0.001, ***

Host-deduced synthetic peptides BiP_Aa_4 and BiP_Aa_6 did not affect planktonic growth even at 112.5 μg/mL, while the presence of BiP_Aa_2, BiP_Aa_5, and BiP_Ml_3 led to some pathogen-specific growth effects. Next, all three synthetic peptides reduced the planktonic growth of *P. aeruginosa* and *S. aureus* by 1.5 – 5 %, while *S. epidermidis* was affected only by BiP_Aa_5 (3 %). The biofilm-preventing potential of all synthetic peptides was tested in various concentrations ranging from 0.4 μg/mL to 112.5 μg/mL against the four opportunistic pathogens using the crystal violet assay (**Fig. 3**). The presence of the control peptide IDR-1018 reduced biofilm formation of *K. oxytoca* at concentrations of 3.5 μg/mL by 30 % and at the highest concentration (112.5 μg/mL) by 70 %. However, no interference with the biofilm formation of the second Gram-negative pathogen, *P. aeruginosa*, was observed. For Gram-positive *S. epidermidis*, biofilm formation was reduced in a range of 10 to 30 %, almost regardless of the concentration, whereas *S. aureus* showed a concentration-dependent reduction in biofilm formation by up to 60 %. For all identified host-derived synthetic peptides, effects on *K. oxytoca* biofilm formation were detected at concentrations above 14 μg/mL, with BiP_Aa_2 and BiP_Aa_5 demonstrating the most substantial effects. In general, the biofilm formation of *P. aeruginosa* was not significantly impacted, but slight interference by BiP_Aa_5 and BiP_Ml_3 was obtained at high concentrations. Depending on the peptide, *S. epidermidis* biofilm formation was reduced by 5 up to 45 %. Notably, the prevention occurred almost regardless of the peptide concentration, which was also the case for the control peptide. Moreover, the effect of BiP_Aa_2 appears to reverse with increasing concentrations. For *S. aureus*, BiP_Aa_2, BiP_Aa_4, and BiP_Ml_3 could not interfere with biofilm formation, whereas the presence of BiP_Aa_5 revealed a reduction of 40 %.

In conclusion, biofilms of Gram-negative and Gram-positive pathogenic bacteria were prevented in a concentration-dependent manner by most of the host-derived synthetic peptides, with BiP_Aa_5 demonstrating the most substantial effects on biofilm formation for all tested pathogens. This finding argues that the initially observed biofilm-preventing activities of the 3 kDa cell extract fractions are mainly based on the respective sequence-identified peptides. Though, other inhibitory biomolecules present in the 3 kDa fractions cannot be excluded entirely.

**Figure 3:**
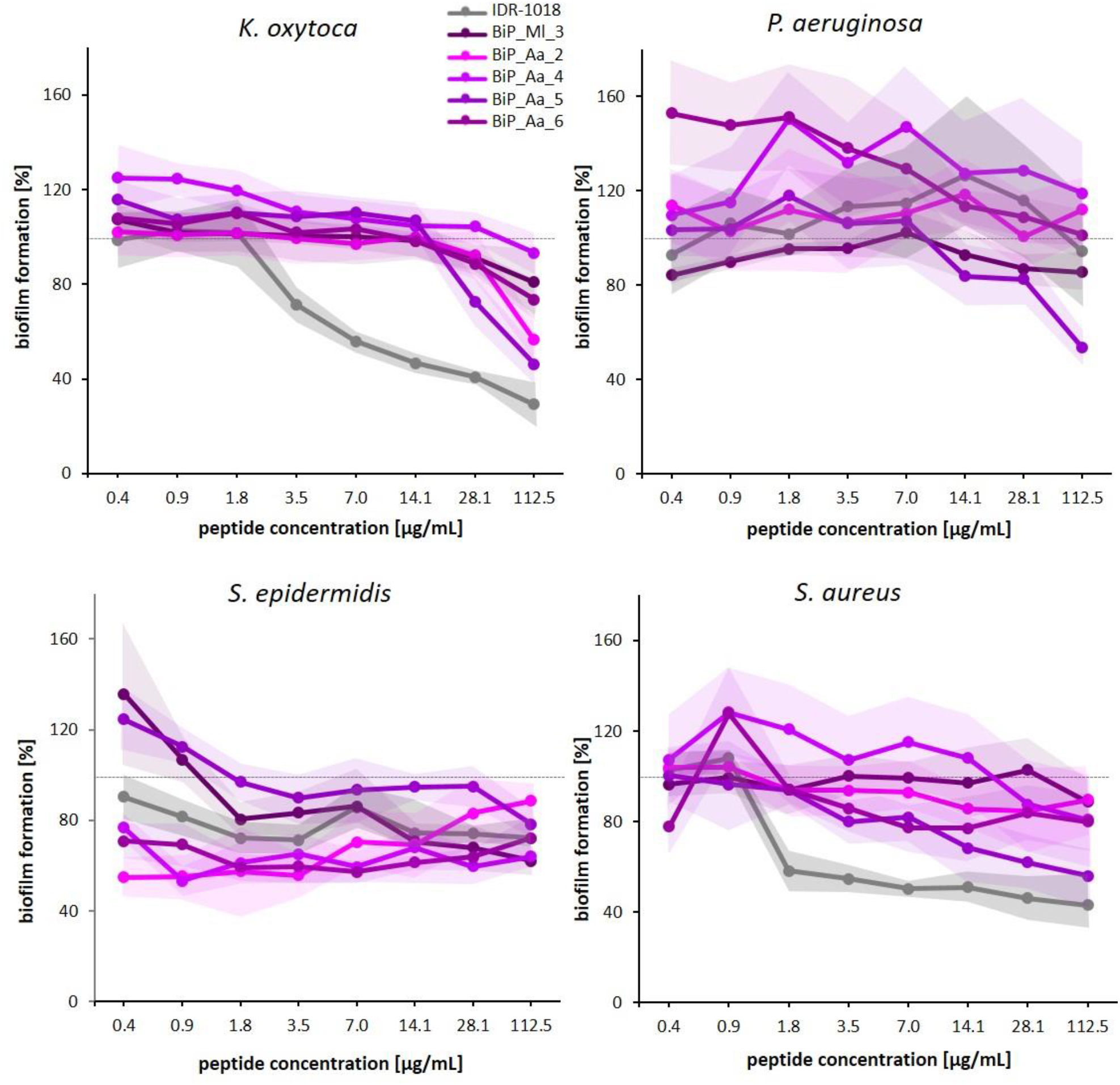
Biofilm-preventing effects of synthetic peptides on static pathogenic biofilms. Biofilm-forming strains *K. oxytoca, P. aeruginosa, S. epidermidis, and S. aureus* (3 × 10^8^ cells/mL) were grown for 18 h in 200 μL Caso Bouillion. Synthetic peptides were added from the beginning in concentrations of 0.4, 0.9, 1.8, 3.5, 7.0, 14.1, 28.1, and 112.5 μg/mL. Diagrams represent the average of three biological, each with eight technical replicates. Biofilm formation was quantified by crystal violet assay. The biofilm-preventing effect was calculated as a percentage value compared to the biofilm control (100 %, dashed line).

### 3.4 Synthetic peptides - biofilm-preventing effects on dynamic *K. oxytoca* biofilms

In the first attempt, microfluidic flow cells were constructed and established for biofilm formation of the opportunistic pathogen *K. oxytoca* (**Fig. 4A**). A comprehensive biofilm formation of *K. oxytoca* was reached with an initial cell concentration of 8 × 10^5^ cells/channel and a flow rate of 15 μl/h for 24 h at 30 °C. A compact biofilm with a wavy surface was formed with a mean biofilm thickness of 11 ± 6 μm (**Fig. 4B** left bar) and volume of 112 ± 62 μm^3^ (**Fig. 4B** left panel, medium control). In a second step, the biofilm formation of *K. oxytoca* was analyzed in the presence of the synthetic host-derived peptides BiP_Aa_2, BiP_Aa_5, and BiP_Aa_6 and the control peptide IDR-1018 (10 ng/channel) with four biological replicates, each with four technical replicates (**Fig. 4B**). In contrast to the medium control, the control peptide IDR-1018 significantly prevented biofilm formation. Predominantly, microcolonies but no 3D structures were detected, resulting in a reduced maximum thickness of 6 ± 2 μm (p = 0.0036) and a reduced volume of 32 ± 16 μm^3^ (p = 0.0046)(**Fig. 4B**). Using the two host-derived synthetic peptides, BiP_Aa_2 and BiP_Aa_5, strong biofilm-preventing effects were observed, confirming the findings for static *K. oxytoca* biofilms. Single cells attached to the surface and formed microcolonies of 8 ± 3 μm (p = 0.1349), resulting in a volume for BiP_Aa_2 of 34 ± 12 μm^3^ (0.0046). Similarly, BiP_Aa_5 formed 7 ± 2 thick biofilms (p = 0.0342) with a volume of 51 ± 25 μm^3^ (p = 0.0309)(**Fig. 4B**).

**Figure 4:**
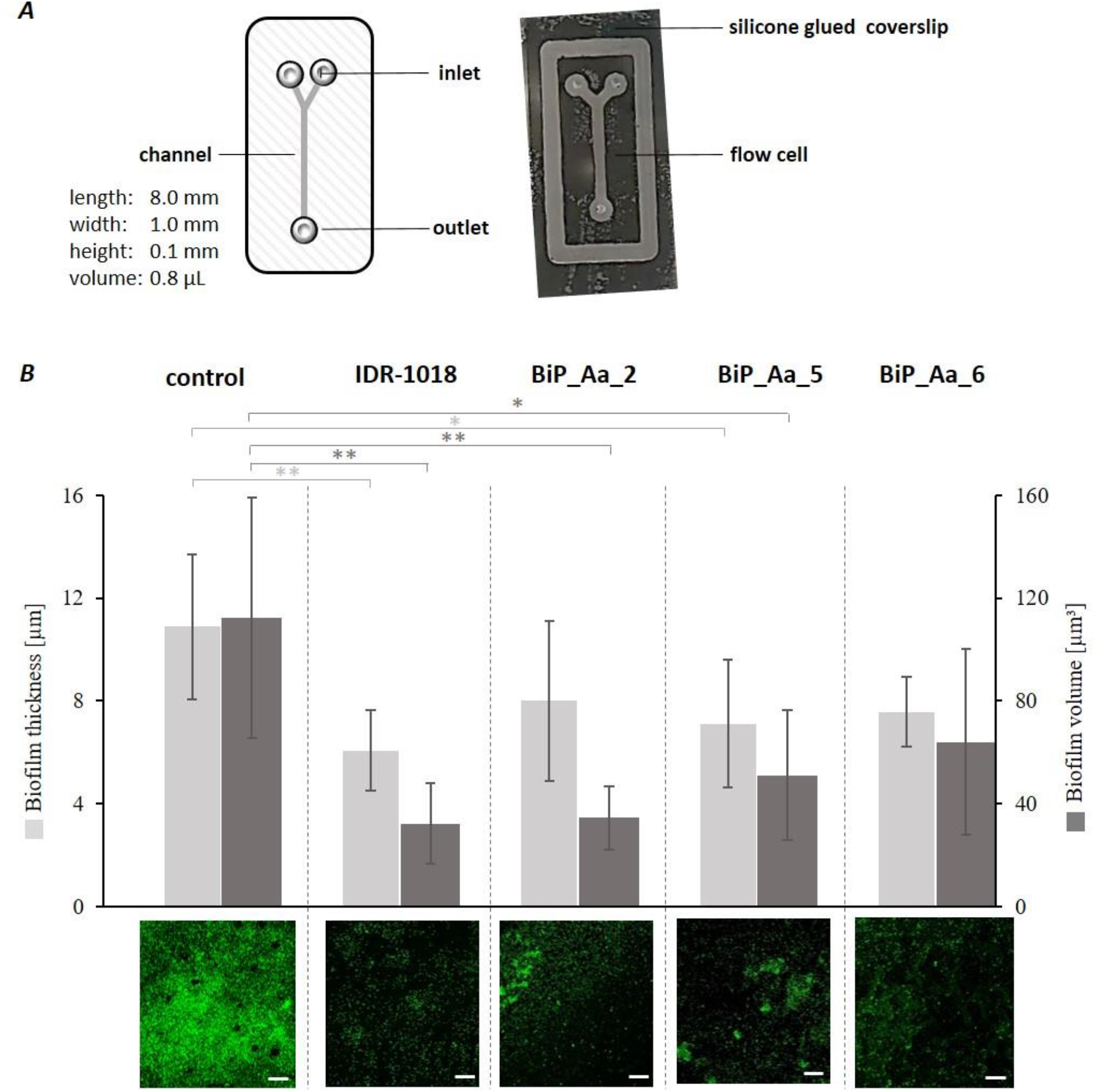
Biofilm prevention of *Klebsiella oxytoca* by selected synthetic peptides in microfluidic flow cells. (***A***) Microfluidic flow cells were constructed of polymethyl methacrylate with a single channel (8 × 1 × 0.1 mm). Silicone adhesive fixed the flow cell to a borosilicate coverslip (24 × 60 × 0.17 mm). (***B***) *K. oxytoca* biofilms were formed in GC minimal media for 24 hours at 30 °C. Synthetic peptides were added in a concentration of 25 μg/mL (10 ng/channel) after 1 h of adhesion time. Peptides and medium were continuously supplemented at a flow rate of 15 μL/h, for 24 h and at 30 °C. Biofilms were stained with SYTO9 and analyzed by confocal laser scanning microscopy using the software Zen black and Imaris. Biofilm characteristics are presented in a bar plot as four biological, each with four technical replicates (upper panel). Significance is represented by the p-values P < 0.05, *; P < 0.01, **. Representative CLSM images with scale bars representing 50 μm (lower panel).

The smallest effect was observed for BiP_Aa_6. Here, the formation of macrocolonies was detected, which reached a thickness of up to 8 ± 1 μm (p = 0.0674) and a volume of 64 ± 36 μm^3^ (p = 0.1443). However, those macrocolonies did not show a compact structure like observed in the medium control (**Fig. 4B**). Consequently, all tested synthetic peptides showed inhibitory effects on dynamic biofilm formation of *K. oxytoca* in flow cells leading to reduced biofilm thickness and volume.

## 4. Discussion

We identified small peptides of 10 – 22 aa length, derived from cDNA expression libraries of two basal marine invertebrates, *A. aurita* and *M. leidyi*, with biofilm-preventing activities (**Fig. 1, 4B**). These marine invertebrates potentially express the peptides as a natural defense, a component of their innate immune system. Examples of such small, cationic, amphipathic peptides are already described for Arthropoda, Mollusca, and Urochordata [20]. A prominent Cnidarian AMP is Aurelin, purified from the mesoglea of *A. aurita* [32]. It is a 40-aa antimicrobial peptide with a molecular mass of 43 kDa with no reported homology to any known antimicrobial peptides. Aurelin has activity against Gram-negative and Gram-positive bacteria due to its structural features of defensins and channel-blocking toxins. Aurelin is processed from an 84 aa pre-proaurelin containing a putative signal peptide [32,33]. Based on our experimental approach to identify new AMPs in this report, it has to be taken into account that newly identified peptides might also be derived from a larger ORF or protein. During sequence analysis of the cloned inserts, we selected the first small ORF in-frame with the coding sequences of the vector backbone. However, the sORFs often did not reflect the complete insert sequence. Most of the small peptides were embedded in larger genes, often encoding for housekeeping genes, like actin and ribosomal proteins (**Tab. S1**). Consequently, some of the the sORFs might be artifically generated based on the experimental design, and encoded peptides identified, do not necessarily possess an ecological antimicrobial activity in the natural system (jellies).

Nevertheless, the biofilm-preventing activities of the five newly identified peptides were observed against *K. oxytoca, P. aeruginosa, S. epidermidis*, and *S. aureus* (**Fig. 3**). The biofilm prevention was assayed parallel to the well-described 12 amino acids cationic AMP (VRLIVAVRIWRR), named IDR-1018 [17,26,27]. This innate defense component is derived from the natural bovine peptide Bactenecin and was reported to show antibiofilm activity against *P. aeruginosa, Escherichia coli, Acinetobacter baumannii, K. pneumoniae*, Methicillin-resistant *S. aureus, Salmonella typhimurium*, and *Burkholderia cenocepacia* [26,27]. The inhibitory activity of IDR-1018 is based on induced dispersal of cells from biofilms at very low peptide concentrations (0.8 μg/mL). The underlying molecular mechanism was shown to involve binding and subsequent degradation of stress-induced second messenger guanosine pentaphosphate ((p)ppGpp) [27]. In contrast, at higher concentrations (10 μg/mL), the peptide caused cell death of the biofilm cells [27]. Besides, weak direct antimicrobial activity of IDR-1018 on planktonic cells was described (MIC 19 μg/mL) [34]. In this report, we verified an antimicrobial effect of IDR-1018 on planktonic bacteria at a high peptide concentration (112.5 μg/mL) (**Fig. 2**), whereas at low concentration no effect was revealed. Further, the reported 50 % biofilm reduction for the pathogens *S. aureus* and *K. oxytoca* was reconfirmed in the here reported experiments. However, in contrast to the original study, *S. epidermidis* biofilm formation was only affected to a minor extent, and *P. aeruginosa* biofilms were almost unaffected. Further, peptides BiP_Aa_4 and BiP_Ml_3 were poorly soluble in hydrophilic solutions, thus, potentially influencing final peptide concentrations for biofilm inhibition assays.

Concentration-dependent biofilm-preventing properties were similarly observed for the jelly-derived peptides against all tested opportunistic pathogens except *P. aeruginosa* **(Fig. 3)**. *P. aeruginosa* biofilm formation was only affected by BiP_Aa_5 and BiP_Ml_3 at high concentrations. A common concern around using AMPs as new antimicrobials is their high susceptibility to enzymatic degradation by proteases [10,35]. *P. aeruginosa* synthesizes many such proteases that are essential virulence factors to interfere with antibacterial defense mechanisms [36], which migth be the reason for our obersvation. In previous reports, *P. aeruginosa* biofilm inhibition was often described as an antimicrobial effect on planktonic cells [37,38]. Our synthetic peptides BiP_Aa_2, BiP_Aa_5, and BiP_Ml_3 showed only low antimicrobial effects on planktonic bacteria (2 – 5 % growth reduction).

A notable exception of the concentration-dependent trend in biofilm prevention against the other opportunistic pathogens was observed for BiP_Aa_2, which showed an increase in the biofilm mass of *S. epderimidis*, when peptide concentration increased above 3.5 μg/mL. Preliminary results demonstrated that an increasing peptide concentration causes dimerization of BiP_Aa_2, likely accompanied by a loss of activity, as shown for the ceratotoxin-like peptide from *Hypsiboas albopunctatus* [39,40]. Overall, peptide BiP_Aa_5 demonstrated the most substantial effects on biofilm formation for all tested pathogens. The reduction of biofilms was comparable to that of IDR-1018, resulting in a reduction of biofilm masses by up to 50 %. This finding agrees with the high similarity of amino acid sequences of the two peptides.

The synthetic peptides, BiP_Aa_2, BiP_Aa_5, and BiP_Aa_6, showed the best effects on the Gram-negative model biofilm former *K. oxytoca*. Consequently, their biofilm-preventing potentials were further validated and studied in spatial resolution in microfluidic flow cells with *K. oxytoca* (**Fig. 4**). Microfluidics technologies can be applied to study bacterial adhesion and biofilm development by precisely controlling the fluidic environment and allows imaging of the biofilm [41]. Microscale systems can address many of their macroscale counterparts’ disadvantages and measurement challenges [42]. Notably, such a system requires relatively fewer sample and reagent volumes, thus advantageous in antibiofilm studies with antimicrobials. The biofilm formation of *K. oxytoca* was analyzed under the constant presence of synthetic peptides after one hour of initial cell adhesion. CLSM micrograph analyses of established biofilms after 24 h in general confirmed inhibitory impacts and showed reduced biofilm thickness and biofilm volume in the presence of all peptides tested (**Fig. 4**). BiP_Aa_2 showed similar biofilm-preventing potential as the control IDR-1018 and thus even more prominent effects as observed for static biofilms. Imaging further enabled for comparison of biofilm structures. *K. oxytoca* generally forms a compact, wavy structure, while adding the peptides revealed that only single cells attached to the surface, successively forming microcolonies. BiP_Aa_6 resulted in the formation of macrocolonies without a compact structure. These findings imply the disturbance of the biofilm matrix, interruption of bacterial cell signaling systems, and disruption or degradation of biofilm-embedded cells’ membrane potential, as reported for AMP actions against bacterial biofilms [43].

Overall, the five characterized invertebrate-derived peptides harbor the typical cationic structure with spatially separated hydrophobic and charged regions resulting in an amphipathic molecule probably crucial for their overall mode of action [44]. The majority showed promising biofilm preventing activities against Gram-negative and Gram-positive opportunistic pathogens and, thus, can serve as starting point to develop efficient and effective antibiofilm agents. Combating bacterial infections caused by biofilm-forming bacteria is a difficult task and a major challenge for healthcare systems and aquaculture [45,46]. AMPs have enormous potential as antibiofilm agents by harboring broad-spectrum antimicrobial activity, a low risk of fast-developing bacterial resistance, and can work synergistically with antibiotics [47]. However, there is still limited information on the interaction of AMPs with biofilm components [43]. More research is needed to understand their precise mechanisms of action on the molecular level, such as inhibiting QS signals that control biofilm formation and interfere with signaling pathways involved in the synthesis of the biofilm matrix [45]. Although dozens of marine invertebrate-derived AMPs are described, their (antibiofilm) mechanisms and applications are still little explored and provide the opportunity for comprehensive research on their use in disease treatment [6]. Our jelly-derived peptides already show promising broad-spectrum antibiofilm activities, and further studies on their mode of action, stability, and side effects can lead to their future application.

## Supplementary Materials

**Figure S1: Identification of biofilm-inhibiting clones from the cDNA expression library using the crystal violet assay**. Cell-free size-fractionated cell extracts were prepared successively from pools of 96, 48, 24, and single clones. Biofilm-inhibiting single clones were further characterized to identify the corresponding peptides.

**Table S1: Identified biofilm-preventing cDNA single clones**. Biofilm-preventing single clones were identified in cDNA expression libraries of *A. aurita* and *M. leidyi* using the crystal violet assay. Activity-conferring sequences were gained and the respective sORFs N-terminally fused to the vector-derived Histidine-tag were translated into peptide sequences. NCBI BLASTp results for those peptide sequences are shown.

## Author Contributions

Conceptualization, R. A. S.; methodology, D. L., L. L., L. G., and N. P.; investigation, D. L., L. L., and L. G.; formal analysis, D. L., L. L., L. G., and N. W.-B.; writing— original draft preparation, N. W.-B., and R. A. S.; writing—review and editing, N. W.-B., and R. A. S.; supervision, N. W.-B. and R. A. S.; project administration, R. A. S.; funding acquisition, R. A. S.. All authors have read and agreed to the published version of the manuscript.

