## Supplementary Material for "Antimicrobial peptides originating from expression libraries of *Aurelia aurita* and *Mnemiopsis leidyi* prevent biofilm formation of opportunistic pathogens"

[2] Current address: Institute of Clinical Molecular Biology (IKMB), Kiel University, Am Botanischen Garten 11, 24118 Kiel, Germany

[*] Joint first authorship

**Tab. S1: Identified biofilm-preventing cDNA single clones.** Biofilm-preventing single clones were identified in cDNA expression libraries of *A. aurita* and *M. leidyi* using the crystal violet assay. Activity-conferring sequences were gained and the respective sORFs N-terminally fused to the vector-derived Histidine-tag were translated into peptide sequences. NCBI BLASTp results for those peptide sequences are shown.

| Single clone designation | Peptide designation | Insert (5'-3') (vector proportion) | Amino acid sequences of the in-frame ORF | NCBI BLASTp | | | |
|---|---|---|---|---|---|---|---|
| | | | | Best homologue | Assession No. | Query Coverage [%] | Identity (%) |
| Aa_112_4H | BiP_Aa_1 | atgcatcatcatcatcatcacatcacaagtttgtacaaaaaagttggcc<br>gccgagtacgatgaaaccgaatctgaagaggaaccatccgaagtagaat<br>cagctgatgatgaaattgaaggtgttgcagcgaataaaggttccgatcc<br>tttattctggcgacgccgaagccgccgccgtattgtctaccgtcgtcgt<br>tggtatacccgtcgtcgttgctactacgctcgccgccgatacatcgctc<br>gccgccgatacatcgctcgcagaagatgctatgctttaataacacgcag<br>aagatggttcagacgaagaggttaacttcccatggcacgaaaattgctg<br>aagacattcgttgattaatcgtagagctgtaaagggtgtttaaaaattg<br>tagtagatgctgattctgattttacgctgataggtgcatgc | RRVR | no significant similarity | | | |
| Aa_112_6C | BiP_Aa_2 | atgcatcatcatcatcatcacatcacaagtttgtacaaaaaagttggcc<br>aactttttttgtacaaagttgtcccccacatag | QLFCTKLSPT | heat shock protein Hsp110 [*Aurelia aurita* ] | AAX09921.2 | 70 | 71 |
| Aa_127_8A | BiP_Aa_3 | atgcatcatcatcatcatcacatcacaagtttgtacaaaaaagttggat<br>ggtcttgagcagtacacagaagtaaaatcagtcatcatcaaaatggacc<br>agaagaatgcttaaagagaccagaacttcaagacaataataaattcatc<br>atgcatatttcatcgttcatcataaatctcgattgtagtttgttgaagg<br>tgccttcttaactgaatgtgtccaactagctaaaaaaaaaaaattcctag<br>ctttttaa | WS | no significant similarity | | | |
| Aa_127_8E | BiP_Aa_4 | atgcatcatcatcatcatcacatcacaagtttgtacaaaaaagttggca<br>cacattctacaatgaacttcgagttgcaccagaggagcatccagtcctg<br>ctcaccgaagctcctttaaatccaaaagctaacagggaaaagatgacac<br>aaattatgttcgaaaccttcaacagccctgcaatgtacgtcgccatcca<br>agccgtactgtccctgtacgcctctggtcgtaccaccggtatcgttctt<br>gattccggagatggtgtcagccacactgtcccaatctacgaaggttatg<br>cccttccccacgccatcatccgtttggatttggctggacgtgatttgac<br>cgactacttgatgaagatcctcaccgagagaggttactcattcaccacc<br>accgccgaaagggaaatcgtcagagacatcaaggagaaactctgctatg<br>tcgcactcgacttccaacaagaaatgctcacagcatcaaccagctcaag<br>cttggagaagagctacgaattacctgacgggacaggtcatcaccatcgg<br>aaacgagagattcaggtgcccagaaacccctcttctaacccgcattcat<br>cggaatggaatcaagcggaatccacgagaccaccatacaaactcaatca<br>tgaaaatgcgatgtcgacatccgtaggacttgcatgccaacaccgtctt<br>tgtctggaggtacgactatgttccccaggtatccgccgacagaatgcac<br>aaggagatcgcttccctcgcacccctcaaccatgaaaattcagatcatc<br>gccccaccagagtaggaaactactcccgtatgggatcggaggctccatc<br>ttggcttcccctctccacctctccaccagatgtcgcatctcgcaatcaa<br>gaatatggatgaatcctgggcccatccattctctcaccacgaaaacgct<br>tccttacaccgcctctgcgcacacttttcaa | THSTMNFELHQRSIQSCSPKLL | retinoid X receptor [*Aurelia aurita* ] | AGT42223.1 | 40 | 71 |
| Aa_127_8F | BiP_Aa_5 | atgcatcatcatcatcatcacatcacaagtttgtacaaaaaagttgggg<br>taaggaacagtgttctattccgtcgcaggcgacttgtatgttagaaagt<br>agtagcactgttttagtagtttaacgataaatcttgaaatgatatctaa<br>caaaatgtgttga | VRNSVLFRRRRLVC | hypothetical protein [*Aurelia aurita* ] | AGN03863.1 | 35 | 100 |
| Aa_127_8H | BiP_Aa_6 | atgcatcatcatcatcatcacatcacaagtttgtacaaaaaagttggga<br>caaactttacaaggaagttcccagttataaactcatcaccccatcagtg<br>gtatctgaaagattgaaggtcagagtttcactcgcacgcacatgcttga<br>aagaactgttgggaaaaggccttatccgggaagtttccaaacatagtga<br>tcagatgatctacactagagctacaaaagaggctgccaaataactttgt<br>ttataagcaatgttgaatcttgaaaatgagtaatctttaaaagcta | TNFTRKFPVINSSPHQWYLKD | toxin TX1 [*Aurelia aurita* ] | AFK76348.1 | 66 | 55 |

| | | | | | | | |
|---|---|---|---|---|---|---|---|
| Ml_068_11H | BiP_Ml_1 | atgcatcatcatcatcatcacatcacaagtttgtacaaaaaagttggaa<br>taatttgagcaataagcgagcttcgatgaaagagagcagaaagaggtag<br>tatcgggggacaaacgaactgtccttaaagcgtaacagactacagaata<br>cacccatcgtgcgttcaggatttttggcaagttcatgtgacagtaaaac<br>gagatgagttcacgagaggtttgatctcaaggagcaactgcgagctata<br>cgacaacgtatacaagtacaaatacgatctccatcaatcagcaccgttc<br>aaatatttggtataaattaatgcagccgggattttgttttgcttccttt<br>ctatttcatagctttgaacaaatacgagaccatggtactcggcgttgtg<br>tactttgattaataataatagtcatagtttgaattatttaaactttcgt<br>agtcttagtctccgcctgtcccattaaagattcttaaatcgctgttgtg<br>actgcgaactttcgcccatttttaacttgtatgcctcaatttttcaaat<br>tcacttcttatgttttaattaatacagttgagaattatgtggctacgag<br>catcagttggagcactgaagctaatatttgttacccggactgtcatact<br>tataggtctcctcatctccataaaaaattaaaaaacaaatggagtactg<br>ttccaattaattttctactattttattgagttttaaaattactttattt<br>actac | II | no significant similarity | | | |
| Ml_011_11H | BiP_Ml_2 | atgcatcatcatcatcatcacatcacaagtttgtacaaaaaagttggca<br>actactattaattaaataatttgaggcaaggctccaggttcttgcgtaa<br>gcggcattcaagatggcgcccaaacacaacaatatggtttttgacagcc<br>atttccacaaggattggcaaaggtatgtcaaaacctggttcaaccaagc<br>cggtaagaagaagcgcaggcgcacaaaacgtatggagaaggctaaagct<br>gtcgcccctcgcccagttgctggtctgctccgacctgttgtacattgtc<br>agactatcagatacaacgcccgagtcagagccggccgaggttttaccct<br>cgacgaactcaaggctgctggtatcaacaagaagcaagctctctcaatc<br>ggaattgctgttgaccacagaaggaagaacaggtctcaggagagtctcc<br>aagctaatgttctcagactgaaggagtacaagagcaggctcatcctctt<br>cccacgaaaggcctccaaacccaagaagggagacagcacccaggctgag<br>atcgatgtcgccactcagctgaccggacctgtcctacccatcaaacaga<br>cctggtctgataccaccgccagagcaatcacggacgaggagaagaaaca<br>atctgctttccagacgatgagaatgtacagagccaatgttcgacttgtt<br>ggagttcgagaaaaacgagccaaggaagccgccgaggatgacggtctgg<br>gagtcaagaagaagaagaaaataatttatttcggtttttttatctttta<br>tcgataatgtca | NYY | no significant similarity | | | |
| Ml_010_9A | BiP_Ml_3 | atgcatcatcatcatcatcacatcacaagtttgtacaaaaaagttggcg<br>gccgcacaactttgtacaagaaagttgggttttttttttttttttaaca<br>tttttaaattttattatttctattttgatttattttatatgacaaatta<br>atttattttcccatttaatctaattgttaaaaaaaaacaaaaaaaaaa<br>ccaaaaacaaaaaaccccccacctttcttgtccaaattggtgatctagg<br>tatattcggatccggttgctaccaaaccccgaaaggaacttgccttggc<br>tgccccccccgctgcccaataactagcataaccccttggggccttcaaa<br>cgggttttgaggggtttttttgctgaaaggaggaacaattcccggttttc<br>ccga | GRTTLYKKVGFFFFF | ND5 gene product [Mnemiopsis leidyi] | YP_004927440.1 | 73 | 100 |
